# A Disease-Associated Mutation Impedes PPIA Through Allosteric Dynamics Modulation

**DOI:** 10.1101/2025.05.09.653011

**Authors:** Yoshikazu Hattori, Munehiro Kumashiro, Hiroyuki Kumeta, Taisei Kyo, Soichiro Kawagoe, Motonori Matsusaki, Tomohide Saio

**Affiliations:** Institute of Advanced Medical Sciences, Tokushima University, Tokushima 770-8503, Japan; Faculty of Advanced Life Science, Hokkaido University, Sapporo, Hokkaido 001-0021, Japan; Student Laboratory, Faculty of Medicine, Tokushima University, Tokushima 770-8503, Japan

## Abstract

Amyotrophic lateral sclerosis (ALS) is a progressive neurodegenerative disease characterized by motor neuron degeneration. Peptidylprolyl *cis-trans* isomerase A (PPIA) is a molecular chaperone involved in protein folding, and its dysfunction has been linked to ALS pathogenesis as proline is recognized as a key residue for maintaining proper folding of ALS-related proteins. A recent study identified a K76E mutation in PPIA in sporadic ALS patients, but its effects on protein function and structure remain unclear. In this study, we used biochemical and biophysical techniques to investigate the structural and functional consequences of the K76E mutation. Our results show that K76E significantly reduces enzyme activity without affecting structure, monodispersity, or substrate recognition. Significant effects of K76E mutation were identified by relaxation dispersion NMR experiments, showing that K76E disrupts key protein dynamics and alters an allosteric network essential for isomerase activity. Corroborated by theoretical kinetic analysis, this dynamics data, revealing the exchange process for K76E approximately one-tenth that of the wild type, explain the reduced *cis-trans* isomerase activity of the K76E mutant. These findings suggest that the pathogenic effect of K76E arises primarily from impaired protein dynamics rather than direct structural disruption. Our study provides new insights into the molecular mechanisms underlying ALS-associated mutations and their impact on protein function.

For Table of Contents

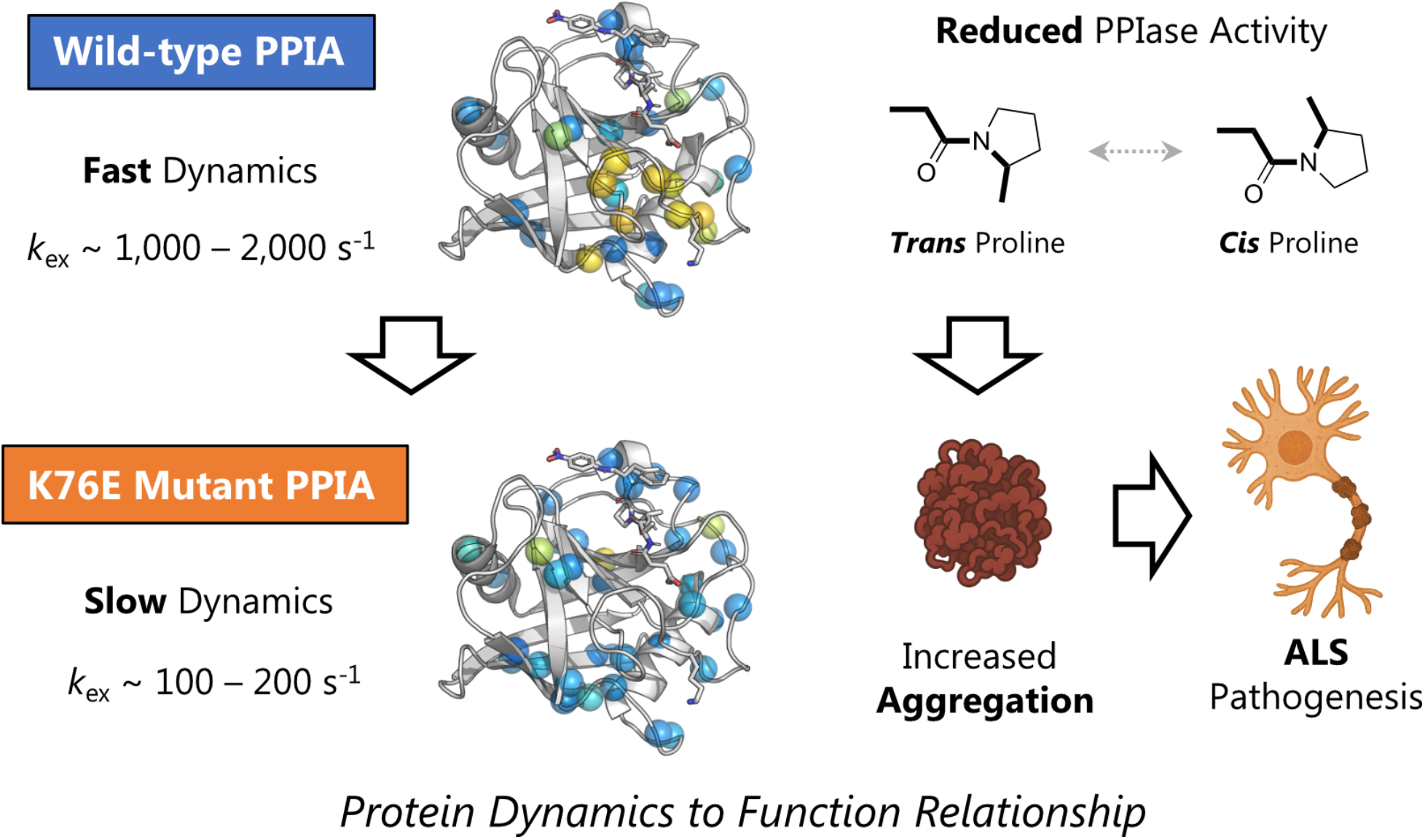

## MAIN TEXT

Amyotrophic lateral sclerosis (ALS) is a fatal neurodegenerative disease caused by motor neuron degeneration, which displays cytoplasmic inclusions.^1,2^ In the processes of ALS pathogenesis, proteins such as transactive response DNA-binding protein 43 (TDP-43), fused in sarcoma, and superoxide dismutase 1 (SOD1) misfold and aggregate, leading to neuronal damage.^3,4^ Moreover, molecular chaperones have gained attention as ALS-associated factors due to their roles in preventing protein misfolding and aggregation.^5,6^ Among them, peptidylprolyl *cis-trans* isomerase A (PPIA, also known as Cyclophilin A), a molecular chaperone possessing peptidylprolyl *cis-trans* isomerase (PPIase) activity,^7,8^ has been proposed as an ALS-related protein.^9^ Indeed, increased levels of TDP-43 and mutant SOD1 aggregates were observed in the spinal cord of PPIA knockout mice, which exhibited accelerated disease onset and progression.^10^ Another study showed that PPIA influenced the polymer structure of heterogeneous nuclear ribonucleoprotein A2 (hnRNPA2), another protein linked to ALS.^11^ Furthermore, proline mutations in TDP-43 and hnRNPA2 caused aggregation by stabilizing self-associated structure.^12^ Thus, PPIA activity is connected to the molecular mechanism of ALS pathogenesis.

More recently, the K76E mutation in PPIA was identified in patients with sporadic ALS.^13^ That study indicated the mutant is less stable and degraded more rapidly in cells than the wild-type protein. Furthermore, molecular dynamics (MD) simulations suggested that K76E mutation induces local structural changes around residues 26–30 and 43–45, which are possibly related to the destabilization of K76E. However, molecular insights are limited and the mechanism of PPIA malfunction caused by K76E mutation remains to be elucidated. More specifically, the impact of the K76E mutation on the PPIase activity of PPIA has not been addressed.

This study focuses on the PPIase activity of PPIA K76E, with biochemical and biophysical measurements. Our biochemical data show that K76E mutation reduces PPIase activity. Despite the distant location of K76 from the substrate-binding site, our biophysical data, particularly from nuclear magnetic resonance (NMR) relaxation measurements, highlight suppression of global conformational dynamics on ms-µs timescale by the K76E mutation. Given the importance of the global conformational dynamics for PPIase activity, the suppression explains the allosteric effect of K76E on the PPIase activity.

We first focused on the impact of the K76E mutation on PPIase activity and performed RNase T1 refolding assay where the slow prolyl *cis-trans* isomerization is the rate-limiting step of the refolding process.^14^ Figure 1A shows the refolding profiles of RNase T1 in the absence and presence of wild-type PPIA and the K76E mutant. In the presence of wild-type PPIA, refolding was strongly accelerated, reflecting its PPIase activity. Refolding was also accelerated in the presence of K76E; however, the acceleration was less pronounced than that observed with wild-type PPIA. This small but significant difference between wild-type PPIA and K76E revealed that the K76E mutation in PPIA reduces the PPIase activity (Figure 1B).

**Figure 1.**
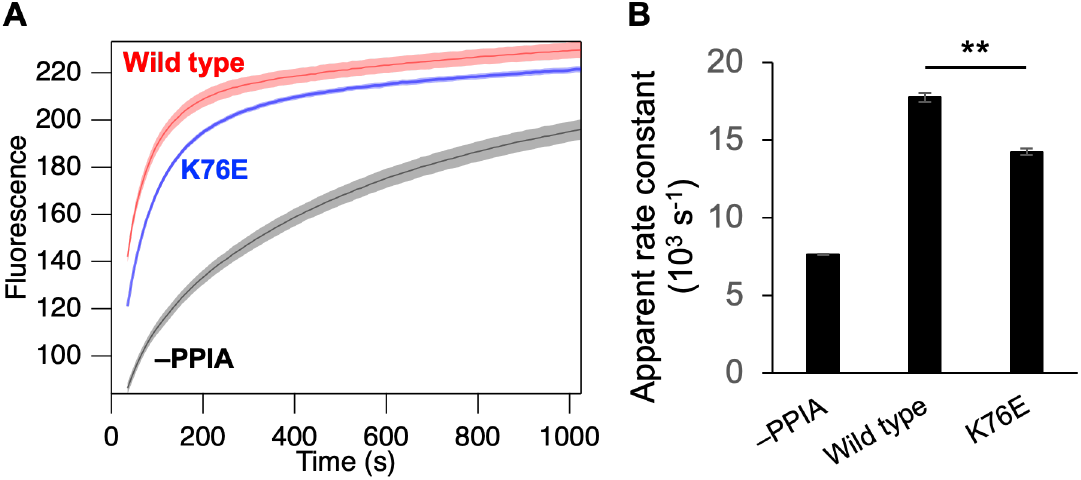
(A) PPIase activity assessed by monitoring the refolding of RNase T1. Fluorescence increases during RNase T1 refolding in the absence of PPIA (black) and in the presence of wild-type PPIA (red) or K76E (blue). Shaded areas represent the standard deviation (N = 3). (B) Apparent rate constants derived from fitting the refolding curves (up to 200 s) to a single exponential function. Statistical significance between wild-type PPIA and K76E was assessed using Welch’s *t*-test. ** indicates *p* < 0.01.

The decrease in PPIase activity was not explained by a decrease in substrate affinity. NMR titration experiments using the model peptides Suc-AAPF-pNA^14^ and Ac-FGPDLPAGD-NH_2_^15^ revealed that the K76E mutation caused negligible change in the affinity (Table 1). The decreased PPIase activity of K76E might potentially be caused by protein aggregation. To investigate aggregation properties, size-exclusion chromatography coupled with multi-angle light scattering (SEC-MALS) analysis was performed. The SEC profiles of wild-type PPIA and K76E both showed sharp peaks with similar elution times. The molecular sizes determined by MALS were 18.2 kDa for wild-type PPIA and 18.3 kDa for K76E, consistent with theoretical value for monomeric PPIA. These data showed that K76E exists as a monodisperse monomer in solution; therefore, the decrease in PPIase activity of K76E is not attributable to differences in aggregation. The decrease in catalytic activity of the K76E mutant might also be attributed to structural changes within the catalytic center. To assess this, chemical shift perturbation (CSP) was analyzed using ^1^H-^15^N correlation NMR spectra measured for wild-type PPIA and the K76E mutant (Figure 2). Overlaying the K76E spectrum onto the wild-type spectrum revealed changes in only a limited number of peaks. CSP mapping showed that most perturbed resonances belonged to residues structurally adjacent to the mutation site. The region near the catalytic center was not significantly perturbed. Thus, the K76E mutant appeared to retain a structure of the catalytic center similar to that of the wild type. Accordingly, we concluded that the reduced activity is unlikely due to structural changes in the catalytic center caused by the mutation.

**Table 1.** Dissociation constants between wild-type PPIA or K76E and substrates.*a*.

|  | Suc-AAPF-pNA | Ac-FGPDLPAGD-NH<sub>2</sub> |
| --- | --- | --- |
| Wild type (mM) | 4.4 ± 0.3 | 0.32 ± 0.1 |
| K76E (mM) | 3.7 ± 0.2 | 0.36 ± 0.1 |

<sup>a</sup> Determined by NMR titration experiments. Errors represent fitting uncertainty.

**Figure 2.**
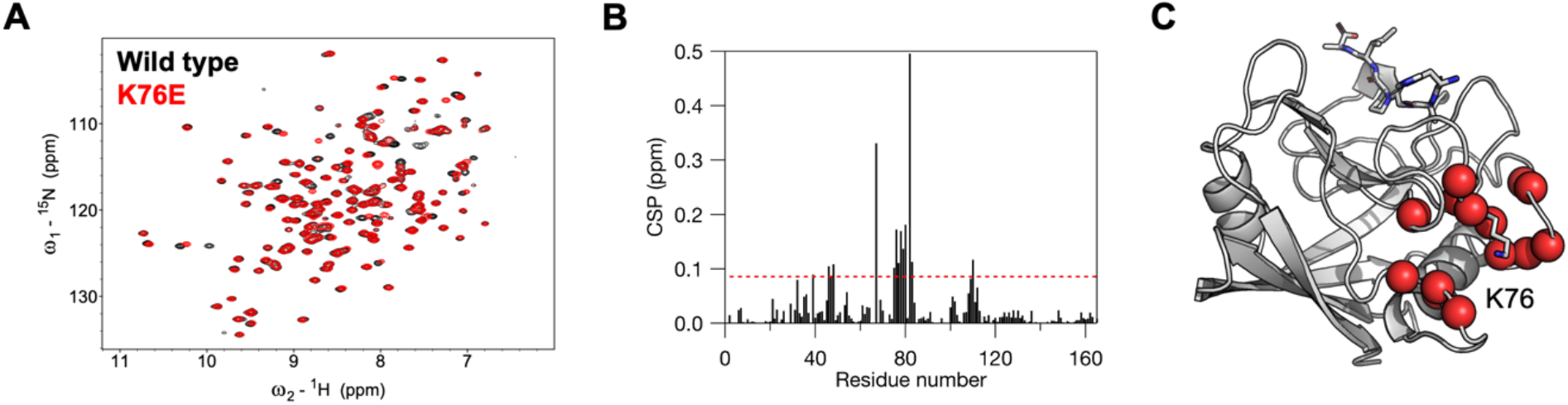
(A) Overlay of 1H-15N correlation spectra of wild-type PPIA (black) and K76E (red). (B) Chemical shift perturbation (CSP) calculated from 1H-15N correlation spectra of wild-type PPIA and K76E. Red dashed line indicates the threshold value (mean + 1 SD). (C) Residues exhibiting CSP above the threshold are mapped as red spheres onto the structure of PPIA (PDB ID: 1RMH). The K76 side chain is shown as sticks.

We also investigated the dynamics of PPIA, given a previous report showing PPIA possesses intrinsic conformational dynamics in a range of 1,000 to 2,000 s^−1^ that are essential for PPIase activity.^16^ To assess the impact of K76E mutation on the global conformational dynamics of PPIA, Carr-Purcell-Meiboom-Gill (CPMG) relaxation dispersion NMR was performed on wild-type PPIA and K76E. For the wild type, several residues located around the substrate-binding site and near K76 showed significant dispersion, indicating relatively fast exchanges processed with rates of a few thousand s^−1^ (Figure 3A and 3B). As proposed in previous studies,^16–18^ these global conformational dynamics drive the *cis-trans* isomerization of the substrate captured by PPIA. In contrast, the relaxation dispersion profiles for residues in PPIA K76E showed obvious change; exchange rates of only a few hundred s^−1^ were observed (Figure 3A and 3B). These results suggest that the enzymatically important exchange processes are attenuated by the K76E mutation. Indeed, prediction of the allosteric network using Ohm^19^ revealed that K76 is part of an allosteric network connected to the catalytic center of PPIA. Thus, the data indicate that the K76E mutation disrupts the allosteric network of PPIA and consequently impedes its functional dynamics.

**Figure 3.**
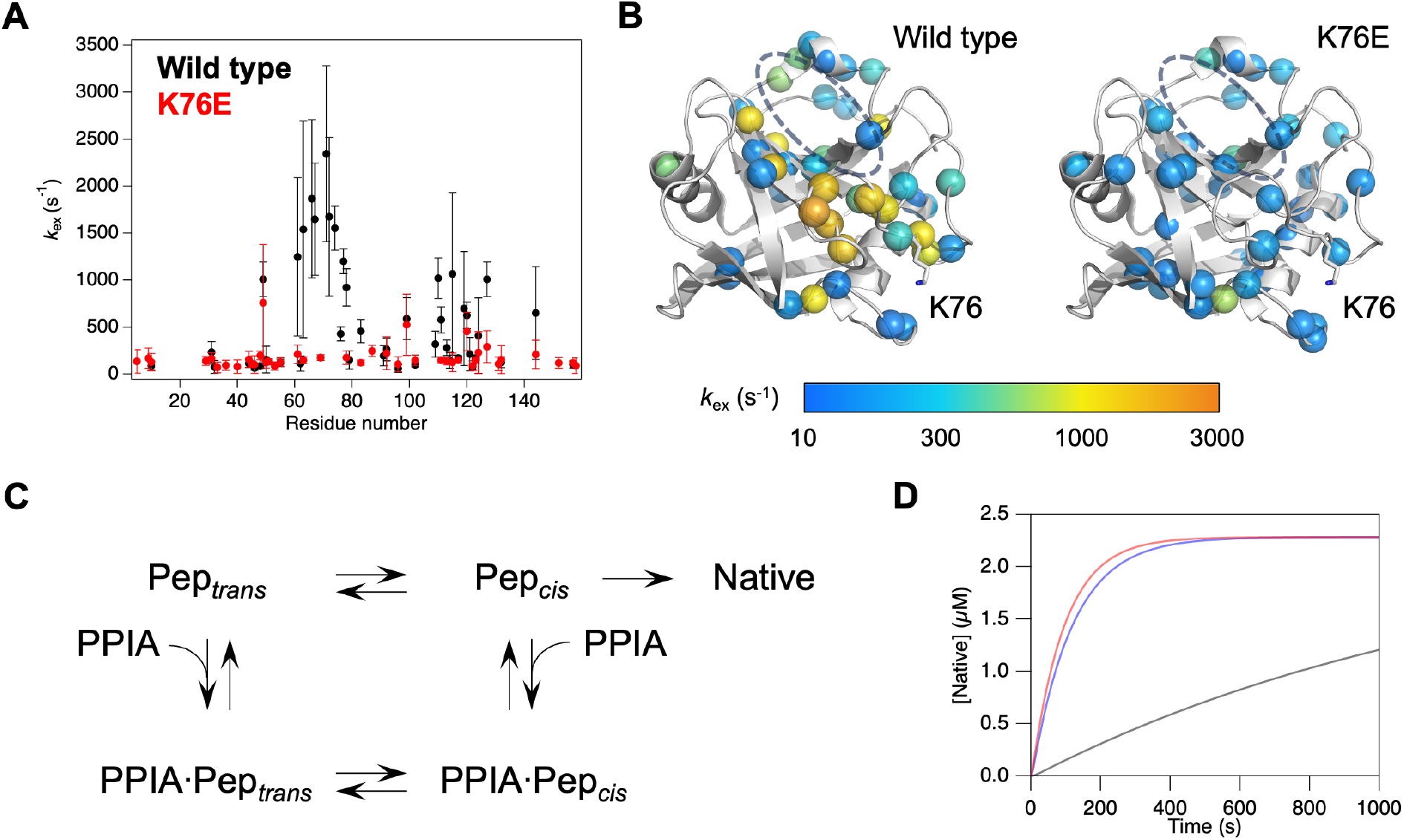
(A) Exchange rate constants (*k*_ex_) for wild-type PPIA (black) and K76E (red) obtained from fitting CPMG relaxation dispersion data. Error bars represent fitting errors. (B) Exchange rate constants (*k*_ex_) mapped onto the structure of PPIA (PDB ID: 1RMH). Residues are shown as spheres colored according to the *k*_ex_ value (gradient bar shown below). The putative substrate binding site is indicated by a grey dashed line. (C) Five-state kinetic model used for simulations of RNase T1 refolding catalyzed by PPIA. (D) Simulated RNase T1 refolding curves in the absence of PPIA (black), presence of wild-type PPIA (red), and presence of K76E (blue), based on the kinetic model in (C).

Given the significant reduction in conformational dynamics, PPIA K76E was expected to exhibit markedly reduced *cis-trans* isomerization activity towards its substrate. This hypothesis was supported by a five-state kinetic model (Figure 3C) reconstituting RNase T1 refolding. Since the refolding processes of RNase T1 are rather complicated, we chose simple model and kinetic parameters following the previous studies.^17,20^ The refolding curves were reasonably simulated if the rates of *cis-trans* isomerization on PPIA K76E was set to approximately one-tenth of those for the wild type (Figure 3D). This indicates that the *cis-trans* isomerization on PPIA K76E was reduced to approximately one-tenth that on the wild type. Therefore, the small but significant reduction in the RNase T1 refolding rate observed for the K76E mutant was explained by drastic reduction of PPIase activity.

Collectively, our biochemical and biophysical data show that the K76E mutation reduces the enzymatic activity of PPIA through the modulation of the dynamics on ms-µs timescale. Previous studies reported the concerted global dynamics of PPIA on ms-µs timescale both in the absence and presence of the substrate and proposed that the dynamics is coupled with *cis-trans* isomerization of the substrate.^16–18,21^ Our study unveiled that a mutation at K76 disrupts the conformational dynamics and PPIase activity of PPIA. Interestingly, K76 is located in one of the two allosteric networks in PPIA, as reported in the previous study,^20^ and the catalytic residue R55 is included in the same network as K76, suggesting that the K76E mutation disrupts the concerted dynamics in the allosteric network. Our data regarding dynamics on the ms-µs scale complement previous results from MD simulations on the order of a few µs.^21^ Although modulations of local fluctuations of PPIA by K76E mutation were probed by MD simulations, the impact of the dynamics modulations to enzyme activity was not understood. In this study, we focused on dynamics on a slower time scale and highlighted a significant effect of K76E to the dynamics on the ms-µs scale that is crucial to the enzymatic activity of PPIA. This allosteric dynamics modulation attenuates the enzymatic activity of PPIA, resulting in a disruption of protein homeostasis, which may have a cumulative effect on the ALS pathogenesis.

## Author Contributions

**Y.H**.: Data curation, formal analysis, funding acquisition, investigation, visualization, writing - original draft, writing - review & editing; **M.K**.: Formal analysis, funding acquisition, investigation, writing - original draft, writing - review & editing; **H.K**.: Investigation, writing - review & editing; **T.K**.: Investigation, writing - review & editing; **S.K**.: Investigation, writing - review & editing; **M.M**.: Funding acquisition, investigation, writing - review & editing; **T.S**.: Conceptualization, funding acquisition, project administration, supervision, writing - review & editing.

## Notes

The authors declare no competing financial interest.

## ACKNOWLEDGMENT

We thank Dr. Eiichiro Mori (Nara Medical University) for critical reading of a manuscript and Dr. Hitoki Nanaura (Nara Medical University) for providing an expression vector coding for PPIA. We thank Dr. Daichi Morimoto (Kyoto University) for providing the program for CPMG data fitting and suggestive comments on analysis. We also thank Azusa Hirano (Tokushima University), and Miyako Sogawa (Tokushima University) for experimental support. This work was supported by Technology Center for Regional R & D, Tokushima University, and the program of the Inter-University Research Network for High Depth Omics, and Joint Usage and Joint Research Programs, the Institute of Advanced Medical Sciences (IAMS), Tokushima University. This work was supported by funding from JSPS KAKENHI (JP23K05657 to Y.H., JP24K18087 to M.K., JP22H04847, JP22K15278, and JP23KK0105 to M.M., JP20H03199, JP20KK0156, JP22H02560, JP22K18361, JP23H05470, JP23H01995, and JP23K23824 to T.S.), HIRAKU-Global Program, which is funded by MEXT’s “Strategic Professional Development Program for Young Researchers” to M.M, MEXT Grant-in-Aid for Transformative Research Areas (B) (JP21H05094 and JP21H05093 to T.S.), AMED (JP22ek0109437, JP25ek0109642, and JP24wm0425004 to T.S.), and JST FOREST Program (JPMJFR204W to T.S.). This work was also partially supported by Astellas Foundation for Research on Metabolic Disorders and Uehara Memorial Foundation to M.M., Takeda Science Foundation Grant, Astellas Foundation for Research on Metabolic Disorders, Nakajima Foundation, Asahi Glass Foundation, Nakabayashi Trust for ALS Research, Kato Memorial Trust for Nambyo Research, Mochida Memorial Foundation for Medical and Pharmaceutical Research, the Naito Foundation, and Serika Fund to T.S.

## METHODS

### Sample Preparation

Human wild-type PPIA (UniProt ID: P62937) and K76E fused with a His_6_-GB1 tag and a TEV cleavage site at the N-terminus were overexpressed in the *E. coli* BL21(DE3) strain. The cells were grown at 37ºC in LB medium or M9 minimal medium containing 1 g/L ^15^NH_4_Cl for unlabeled or ^15^N-labeled samples, respectively. Protein expression was induced with 0.5 mM IPTG at an OD_600_ of 0.6-0.8, followed by cultivation at 15ºC overnight. The cells were harvested and resuspended in lysis buffer containing 50 mM Tris, pH 8.0, and 500 mM NaCl. The suspended cells were lysed using an ultrasonic homogenizer and centrifuged at 18,000 rpm for 30 min. The protein in the supernatant was purified using Ni-NTA agarose (QIAGEN), and TEV protease was added to the eluate during dialysis against 50 mM Tris, pH 8.0, and 150 mM NaCl. After digestion, the solution was purified again using Ni-NTA agarose to remove the His_6_-GB1 tag and any uncleaved protein. The flow-through was further purified by size-exclusion chromatography using a HiLoad 26/600

Superdex 200 pg column (Cytiva) equilibrated with 20 mM potassium phosphate, pH 7.0, 100 mM KCl, 4 mM 2-mercaptoethanol, and 0.05% NaN_3_.

RNase T1 and Suc-AAPF-pNA were purchased from Sigma-Aldrich. Ac-FGPDLPAGD-NH_2_ was synthesized by GenScript.

### RNase T1 Refolding Assay

RNase T1 refolding assay was performed as described previously.^14^ Briefly, RNase T1 was denatured by incubating overnight with 6.9 M urea. Urea-denatured RNase T1 (12 μL) was rapidly diluted by manual pipetting 15 times to 408 μL of pre-incubated buffer or PPIA solution (100 mM Tris, pH 8.0; dilution rate: 35-fold) to initiate refolding of RNase T1. The dead time was 20 s. The time-course of tryptophan fluorescence of 2.28 μM RNase T1 in the presence and absence of 0.05 μM PPIA (wild type and K76E) was monitored using a spectrofluorometer (FP-8350; JASCO). The excitation and emission wavelengths were set to 280 nm (bandwidth: 2.5 nm) and 320 nm (bandwidth: 10 nm), respectively. Each profile represents the average of three replicates recorded at a scan speed of 10 s and response time of 8 s. The standard error was estimated from the three replicates. All the measurements were performed at 10°C. The baseline (buffer- or PPIA-only control) was subtracted. The photobleaching was suppressed by placing a 1% dimmer over the excitation light window.

### Statistical Analysis

Statistical analysis was performed using Welch’s *t*-test to compare initial rate constants between wild-type PPIA and K76E. Analysis was conducted in R (version 4.4.1). A p-value less than 0.05 was considered statistically significant. Sample sizes for each group are indicated in the figure legend.

### SEC-MALS

SEC-MALS was measured using DAWN HELEOS8+ (Wyatt Technology Corporation, Santa Barbara, CA, USA), a high-performance liquid chromatography pump LC-20AD (Shimadzu, Kyoto, Japan), refractive index detector RID-20A (Shimadzu), and UV-vis detector SPD-20A (Shimadzu), located downstream of the Shimadzu liquid chromatography system and connected to a PROTEIN KW-803 gel filtration column (Shodex). Differential RI (Shimadzu) downstream of MALS was used to determine protein concentrations. The running buffer comprised 20 mM potassium phosphate, pH 7.0, 100 mM KCl, 4 mM 2-mercaptoethanol, and 0.05% NaN_3_. The 100 µL of 50 μM wild-type PPIA or K76E was injected at a flow rate of 1.0 mL min^−1^. The data were analyzed using ASTRA version 7.0.1 (Wyatt Technology Corporation). Molar mass analysis was performed over half the width of the top height of the UV peak.

### NMR

For the measurements of ^1^H-^15^N correlation spectra of wild-type PPIA and K76E, the samples were dissolved in 20 mM potassium phosphate, pH 7.0, 100 mM KCl, 4 mM 2-mercaptoethanol, and 0.05% NaN_3_ containing 5% D_2_O at a protein concentration of 0.1 mM. The measurements were performed on a Bruker AVANCE III 500 MHz spectrometer equipped with a BBFO probe at 25ºC. Backbone resonance assignments of wild-type PPIA were obtained from BMRB entry 27265. The resonances of K76E were assigned mainly by tracing the resonances of the wild type and were additionally confirmed by using triple resonance data, including HNCA, HNCOCA, HNCACB, and CBCACONH, which were acquired with a Bruker AVANCE III HD 600 MHz spectrometer equipped with a TBI probe at 25ºC. The data were processed using NMRPipe.^23^ The spectra were analyzed using POKY.^22^ The weighted average of the chemical shift perturbation (CSP) was calculated using following equation:

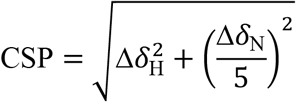

where Δδ_H_ and Δδ_N_ are the ^1^H and ^15^N chemical shift changes, respectively.

For the titration experiments with substrates, the samples were dissolved in 50 mM sodium phosphate, pH 6.5, and 1 mM dithiothreitol. NMR spectra were measured on a Bruker AVANCE III 500 MHz spectrometer equipped with a BBFO probe at 25ºC. The titration with Suc-AAPF-pNA was performed at a 0.5 mM protein concentration. A 200 mM stock solution of Suc-AAPF-pNA in dimethylsulfoxide-d_6_ was added to the protein solution to achieve final molar ratios of 2, 4, 8, and 16. Samples were prepared separately to maintain a final concentration of 4% dimethylsulfoxide-d_6_. The titration with Ac-FGPDLPAGD-NH_2_ was performed at a 0.2 mM protein concentration. A 20 mM stock solution of Ac-FGPDLPAGD-NH_2_ in buffer was added sequentially to the protein solution to achieve final molar ratios of 0.5, 1, 2, 4, and 8. The data were processed using NMRPipe.^23^ The spectra were analyzed using POKY.^24^ Dissociation constants (*K*_d_) were determined by fitting the observed chemical shift changes (Δ_obs_) to the standard quadratic binding equation for a 1:1 interaction:^22^

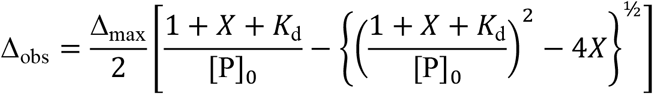

where *X* is the molar ratio of substrate to protein, [P]_0_ is the initial concentration of protein, Δ_max_ is the chemical shift changes between the fully bound and unbound forms, and Δ_obs_ is the chemical shift changes between at a given titration point relative to the unbound form. In the case of the titration with Suc-AAPF-pNA, ^1^H chemical shift changes from five residues were globally fitted with the equation because ^1^H shift changes were dominant. In the case of the titration with Ac-FGPDLPAGD-NH_2_, the weighted averaged chemical shift changes (calculated using the CSP equation above) for 10 residues were globally fitted using the binding equation.

For the measurements of ^15^N TROSY-based CPMG relaxation dispersion, wild-type PPIA and K76E were prepared at a 0.7 mM concentration in 50 mM sodium phosphate, pH 6.5, and 1 mM dithiothreitol. NMR spectra were measured on a Bruker AVANCE Neo 800 MHz spectrometer equipped with a cryogenic TCI probe at 10ºC. CPMG pulses were applied at frequencies (*ν*_CPMG_) of 31.25, 62.5, 125, 250, 375, 500, 625, 750, and 1000 Hz during a 16 ms mixing period. The data were processed using NMRPipe.^25^ Two-site exchange rate constants (*k*_ex_) were extracted using GLOVE^26^ by the fitting the data to the Luz-Meiboom equation:^27^

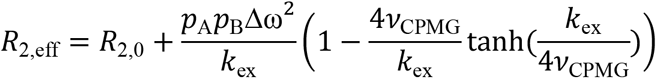

where *R*_2,eff_ is the effective transverse relaxation rate, *R*_2,0_ is the transverse relaxation rate in the absence of chemical exchange, *p*_A_ and *p*_B_ are the populations of the states in the two-state model, Δ_ω_ is the chemical shift difference between the states, and *ν*_CPMG_ is the frequency of the CPMG pulses.

### Mapping Allosteric Communications

Allosteric coupling intensities within PPIA residues were predicted for the atomic coordinate of PDB ID: 1RMH using Ohm web-server.^19^ An active site was designated to R55. The distance cutoff is set to 3.4 Å, and the parameter ‘alpha’ is set to 3.

### Theoretical Kinetic Analysis

Protein folding in which proline *cis-trans* isomerization is the rate-limiting step was simulated in the presence and absence of PPIA (wild type and K76E) using Kintek Explorer (v.11.1.1; Kintek Corporation).^8^ The mechanism tested here was built upon the known mechanism for minimal reaction model of PPIA isomerization,^17^ which consisted of four states and eight reaction pathways, as shown in Figure 3C. The microscopic rate constants were set based on previous studies.^17,20^ Based on the NMR results, the rate constants for the transition between PPIA:Pep_*trans*_ and PPIA:Pep_*cis*_ for PPIA K76E were fixed to approximately one-tenth of those for the wild type. To account for the folding pathway after isomerization, a pathway from Pep_*cis*_ to Native states was included in the model. The rate constant for the Pep_*cis*_-Native transition was set to 0.1 s^−1^, which roughly reproduces the refolding behavior of RNase T1 in the presence and absence of PPIA (Figure 3D). Differential equation of the kinetic model was numerically solved under the initial conditions of 2.28 μM Pep_*trans*_, 0 or 0.05 μM PPIA, and 0 μM other components.

